# *Osmia bicornis* is rarely an adequate regulatory surrogate species. Comparing its acute sensitivity towards multiple insecticides with regulatory *Apis mellifera* endpoints

**DOI:** 10.1101/366237

**Authors:** Philipp Uhl, Osarobo Awanbor, Robert S. Schulz, Carsten A. Brühl

**Affiliations:** University of Koblenz-Landau, Institute for Environmental Sciences, Fortstrasse 7, 76829 Landau, Germany

## Abstract

Bee species provide essential ecosystem services and maintain floral biodiversity. However, there is an ongoing decline of wild and domesticated bee species. Since agricultural pesticide use is a key driver of this process, there is a need for a protective risk assessment. To achieve a more protective registration process, two wild bee species, *Osmia bicornis* and *Bombus terrestris*, were proposed by the European Food Safety Authority as additional test surrogates. We investigated the acute toxicity (median lethal dose, LD50) of multiple commercial insecticide formulations towards the red mason bee (*O. bicornis*) and compared these values to honey bee (*Apis mellifera*) regulatory endpoints. In two thirds of all cases *O. bicornis* was less sensitive than the honey bee. By applying an assessment factor of 10 on the honey bee endpoint a protective level was achieved for 87% (13 out 15) of all evaluated products. Our results show that *O. bicornis* as a non-sensitive species is rarely an adequate additional surrogate species for lower tier risk assessment. Given the currently limited database, the honey bee seems sufficiently protective in acute scenarios as long as a reasonable assessment factor is applied. However, additional surrogate species such as *O. bicornis* and *B. terrestris* are still relevant for ecologically meaningful higher tier studies.

## Introduction

Bees are important pollinators of wild and cultivated flora which makes them essential providers of ecosystem services and maintainers of floral biodiversity [1, 2]. Aside from the honey bee *Apis mellifera* there is a broad spectrum of wild bee species that contribute substantially to plant pollination [3]. However, there is an ongoing trend of wild bee species decreasing in abundance and diversity all over the world. Furthermore, honey bee hive numbers are also substantially decreasing in North America and many European countries [4]. Among various environmental factors, e.g. habitat loss & fragmentation, parasites, agricultural pesticide use has been identified as one the key drives of bee decline [5]. The ecological challenge of flying insect decline in general seems to have been underestimated and consequently disregarded in the past: As a recent study by Hallmann et al. (2017) shows, there has been a severe 75% decline in flying insect biomass in several German natural reserves over roughly the last three decades [6].

In the European agricultural landscape, bees are exposed to a variety of pesticides that target all major pests, e.g. herbicides, fungicides, insecticides [7, 8]. They are not only contaminated during foraging on crops but also from visitations of field-adjacent wild flowers [9]. Bees are exposed to pesticides by direct overspray as well as oral uptake of and contact to nectar and pollen while foraging. There are also fed contaminated pollen and nectar as larvae. Furthermore, there is potential uptake of soil residues by adults and larvae of soil-nesting species [10]. Moreover, consumption of non-nectar fluids such as puddle water, guttation droplets or extrafloral nectar may also lead to contamination [11, 12]. Consequently, bee species are exposed to pesticides through various environmental matrices throughout their lifespan.

To prevent adverse impact of pesticide applications on wild bee populations, toxic effects of these substances on bee species need to be understood. However, the majority of toxicity testing in laboratory and field setups has been performed using the honey bee, a bred livestock species, whereas all other bee species are far less well-understood in their sensitivity [10].

Furthermore, the honey bee is the only pollinator species that is tested for its reaction towards pesticides in the current risk assessment scheme after Regulation (EC) 1107/2009 [13]. However, wild bee species (i.e. bumble bees, solitary bees) may show quite different responses to pesticide exposure due to differences in physiology and ecology [14]. As a reaction to the information scarcity regarding the sensitivity of bumble bees and solitary bees, the European Food Safety Authority (EFSA) proposed the inclusion of the buff-tailed bumblebee *Bombus terrestris* and the red mason bee *Osmia bicornis* into EU pesticide risk assessment as additional surrogate species [15]. However, there has been reasonable doubt that these two species are adequate to provide additional safety in lower tier risk assessment. Uhl et al. (2016) tested five European bee species in acute contact exposure scenarios with a formulated insecticide product (PERFEKTHION®) containing the toxic standard dimethoate [16]. They found that *B. terrestris* and *O. bicornis* were the least sensitive species when compared to a dataset of their own results and collected literature data. Another study by Heard et al. (2017) compared the acute oral sensitivity of the honey bee towards several pesticides (active ingredients) to *B. terrestris* and *O. bicornis* [17]. They found contrasting sensitivity ratios depending on substance since both wild bee species were sometimes more and sometimes less sensitive. *Bombus terrestris* was generally less sensitive than the honey bee in acute toxicity studies that were compiled by Arena & Sgolastra (2014) [14]. They could not collect *O. bicornis* data but other *Osmia* species (*O. cornifrons*, *O. lignaria*) were usually also more resistant than *A. mellifera.* Moreover, EFSA (2013) proposed an assessment factor of 10 to account for interspecific differences when testing only honey bees [15]. This approach proofed to be protective in 95% of cases in the meta-analysis by Arena & Sgolastra (2014) [14]. It is unclear, however, if this factor would be protective for both proposed test species due to the slim database of their sensitivity [16, 17].

There is a need to assess the suitability of the new test species that EFSA proposed. Only sensitive species will reduce uncertainty in lower tier risk assessment. However, with the current database it is not possible to properly evaluate if the proposed species are adequate. Therefore, we tested one of these two species, *O. bicornis*, with commercial formulations of multiple common insecticides. We performed acute contact toxicity laboratory tests to derive 48h contact median lethal doses (LD50s). We wanted to assess the acute toxic potency of several insecticides from various classes on *O. bicornis*. Furthermore, our goal was to compare those toxicity endpoints to honey bee data from pesticide regulation. This enabled us to evaluate if *O. bicornis* is usually more sensitive than the honey bee which would make it a suitable additional surrogate species. Additionally, we examined if an assessment factor of 10 is protective when comparing honey bee to *O. bicornis* sensitivity.

## Materials and methods

### Insecticides

The majority of tested insecticides were chosen with respect to the application frequency of their commercial products in apple, grapes and winter oilseed rape (Table 1) which represent three main cultivation types in Germany [18]. Additionally, formulations of four insecticides that are not frequently applied were included because of the following reasons: Imidacloprid has been implicated as a major factor in bee decline [10]. Dimethoate is often used as a toxic reference in bee ecotoxicity studies. Chlorpyrifos was chosen to include another organophosphate insecticide aside from dimethoate. Furthermore, flupyradifurone is a relatively new insecticide with low acute toxicity towards honey bees that has been applied for registration in multiple EU countries [19]. Insecticides were assigned to pesticide classes according to the Compendium of Pesticide Common Names [20]. Representative formulated products that contain those pesticides as active ingredients (a.i.) were chosen for testing (Table 1). Most of these formulations are or were registered in Germany in recent years aside from Pyrinex® (a.i. chlorpyrifos) and Sivanto® SL 200 G (a.i. flupyradifurone). To ease readability, only active ingredient instead of formulated product names are used hereafter. Please see Table 1 for a list of all tested formulated products and corresponding active ingredients.

**Table 1.**
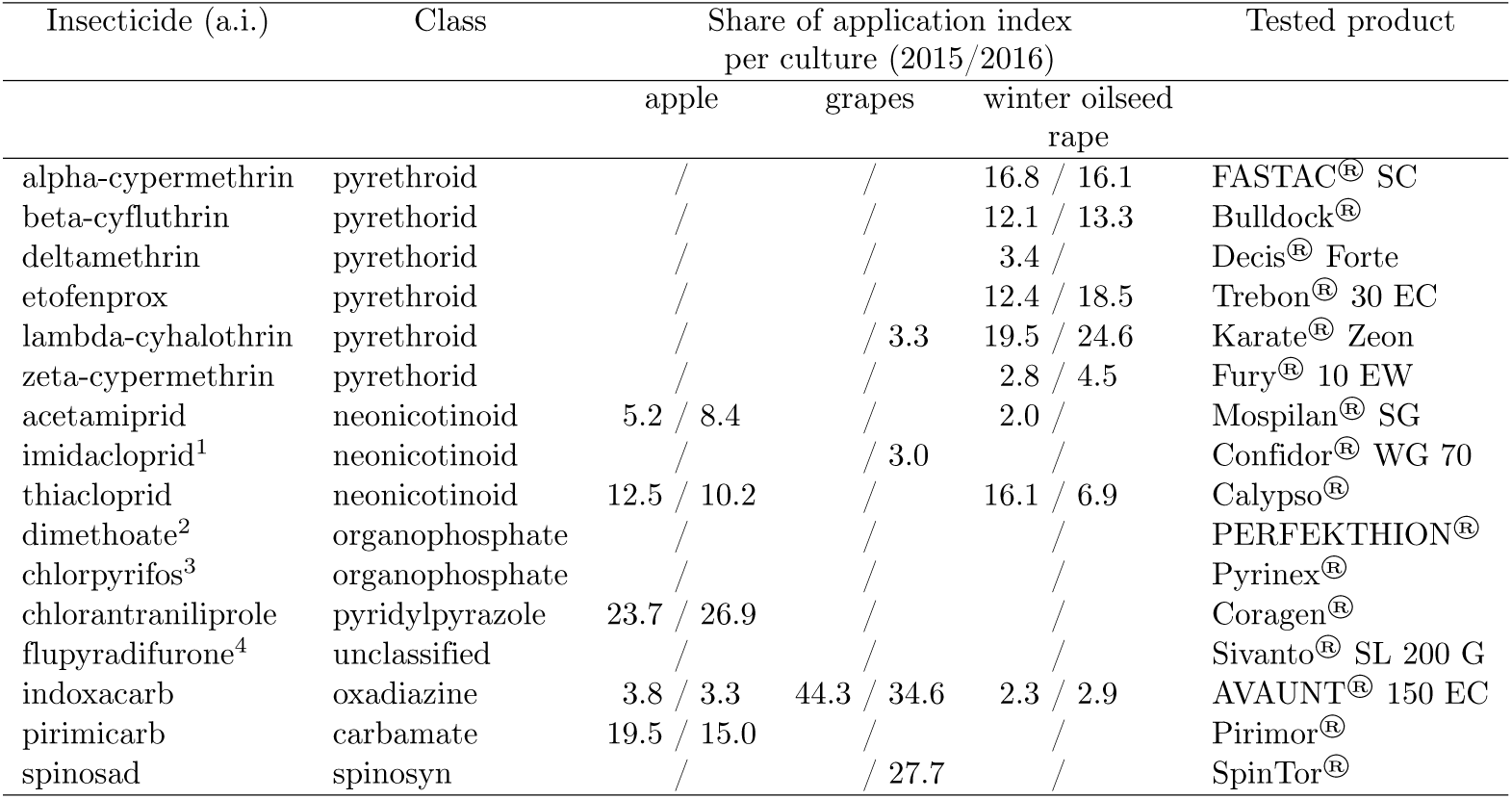
Tested insecticides and their usage shares in German agriculture. The application index is defined as the number of pesticide applications in a crop in relation to the application rate and cultivated area. Data from Julius Kühn-Institut (2018) [18].

| Insecticide (a.i.) | Class | Share of application index<br>per culture (2015/2016) |  |  | Tested product |
| --- | --- | --- | --- | --- | --- |
|  |  | apple | grapes | winter oilseed<br>rape |  |
| alpha-cypermethrin | pyrethroid | / | / | 16.8 / 16.1 | FASTAC® SC |
| beta-cyfluthrin | pyrethroid | / | / | 12.1 / 13.3 | Bulldock® |
| deltamethrin | pyrethroid | / | / | 3.4 / | Decis® Forte |
| etofenprox | pyrethroid | / | / | 12.4 / 18.5 | Trebon® 30 EC |
| lambda-cyhalothrin | pyrethroid | / | / 3.3 | 19.5 / 24.6 | Karate® Zeon |
| zeta-cypermethrin | pyrethroid | / | / | 2.8 / 4.5 | Fury® 10 EW |
| acetamiprid | neonicotinoid | 5.2 / 8.4 | / | 2.0 / | Mospilan® SG |
| imidacloprid<sup>1</sup> | neonicotinoid | / | / 3.0 | / | Confidor® WG 70 |
| thiacloprid | neonicotinoid | 12.5 / 10.2 | / | 16.1 / 6.9 | Calypso® |
| dimethoate<sup>2</sup> | organophosphate | / | / | / | PERFEKTHION® |
| chlorpyrifos<sup>3</sup> | organophosphate | / | / | / | Pyrinex® |
| chlorantraniliprole | pyridylpyrazole | 23.7 / 26.9 | / | / | Coragen® |
| flupyradifurone<sup>4</sup> | unclassified | / | / | / | Sivanto® SL 200 G |
| indoxacarb | oxadiazine | 3.8 / 3.3 | 44.3 / 34.6 | 2.3 / 2.9 | AVAUNT® 150 EC |
| pirimicarb | carbamate | 19.5 / 15.0 | / | / | Pirimor® |
| spinosad | spinosyn | / | / 27.7 | / | SpinTor® |

### Provision of test species

The red mason bee *Osmia bicornis* (LINNEAUS, 1758) was used as test species. Bees were ordered as cocoons (WAB-Mauerbienenzucht, Konstanz, Germany) and stored at 4°C until experimental preparation started.

### Experimental Procedure

Acute, contact toxicity of 16 insecticide formulation towards *O. bicornis* females was investigated (see Supporting Information Table S1 Table for a timeline of the experiments). To that end, a protocol for solitary bee acute contact toxicity testing from the International Commission on Plant-Pollinator Relationships (ICPPR) was followed or partly adapted [21]. This protocol is a precursor of a standardised testing guideline. Prior to the experiments, bee cocoons were collected from the refrigerator and placed in an environmental chamber at test conditions which are explained below to hatch. Male bees were also collected but only used for range finding tests. Female bees’ eclosion time was usually between five to seven days. Afterwards hatched females were again stored at 4°C until one day before application. At this date, they were transferred in to the environmental chamber in test cages (1 L plastic boxes sealed with a perforated lid) and fed *ad libitum* with sucrose solution 50% (w/w) through 2 mL plastic syringes to acclimatize overnight. Twenty bees were assigned to each treatment (usually 5 per cage, n = 4). Please see the raw data for details on individual study setups [22]. Environmental conditions were set to 16:8h day/night rhythm, 60% relative humidity and 21°C. In the summer of 2017 there was a malfunction of the environmental chamber which caused the light to stay on throughout the whole day. Two test runs were therefore conducted with constant lighting (dimethoate, indoxacarb). Since control mortality was below the quality criterium of 10% in those runs, they were evaluated as valid, nonetheless. Anaesthetisation of bees was necessary before the transfer to test cages. To achieve a calm state, bees were chilled at 4°C. During this process they were also weighed. Bees were anaesthetised a second time before application which was performed in a petri dish. In cases where the ambient temperature was too high to keep bees calm after chilling, petri dishes were put on ice for additional cooling. Moribund bees were rejected and replaced with healthy bees prior to the test start.

Treatment solutions were prepared as follows: a control of deionized water containing 0.5% (v/v) wetting agent (Triton™ X-100, Sigma-Aldrich) and at least five insecticide treatment solutions. Concentrations and number of insecticide treatments were determined after conducting range finding tests with male bees before the main test. Results of these pretests were extrapolated to females using the weight difference of both sexes. Insecticide solutions were prepared by diluting the respective concentration in deionized water containing 0.5% wetting agent. In the first tests, bees were applied with 2 *μ*L on the dorsal side of the thorax between the neck and wing base using a Hamilton micro syringe (Hamilton Bonaduz AG). Due to easier handling, an Eppendorf Multipette® plus (Eppendorf AG) was used later on for most of the tests. In three tests (chlorantraniliprole, flupyradifurone, pirimicarb) the applied volume had to be increased to 4 *μ*L to dilute high doses. Please see the raw data published in Uhl et al. (2018) for details [22]. After ten to 15 min the treatment solution was fully absorbed and a paper tissue was inserted into test cages to provide a hiding place. Following the application bees were returned to the environmental chamber and fed 50% sucrose solution *ad libitum.* Mortality was assessed after 24, 48, 72 and 96h. For dimethoate a second test run was performed as part of an ICPPR ringtest. Control mortality was ≤10% in all experiments except for flupyradifurone and chlorantraniliprole (both 15%). Those two cases were evaluated and are considered valid since in the ICCPR test protocol it is discussed that control mortality thresholds might be increased to 15 or 20% in the long run.

### Data analysis

Median lethal dose values (contact 48h LD50) were calculated for all tested insecticidal products by fitting a dose-response model to the data. Please see Uhl et al. (2018) for an account of the raw data [22]. Models were chosen by visual data inspection and using Akaike information criterion (AIC). Furthermore, it was ensured that appropriate model were used for tests with control mortality (no fixed lower limit). Where multiple LD50 values were available a geometric mean LD50 was computed. Weight-normalised LD50 values were further calculated by dividing LD50 values by mean fresh weight of all bees in a respective test. All statistical analyses were conducted with R 3.4.4 [23]. We used the “drc” package [24] for dose-response modeling (version 3.0-1). Honey bee contact 48h LD50 values were gathered by screening regulatory documents (EC review, report, EFSA conclusion, rapporteur member state draft/renewal assessment reports). Furthermore, we contacted national and European authorities, manufacturers and EFSA to collect data and verify them. For a detailed account of the data collection process and various data sources please see Supporting Information Appendix S1 Appendix and Tables S2 Table, S3 Table. To compare *A. mellifera* and *O. bicornis* endpoints sensitivity ratios (R = LD50_*A. mellifera*_ / LD50_*O.bicomis*_) were calculated according to Arena & Sgolastra (2014) for all tested insecticides [14].

## Results

Sensitivity of *O. bicornis* towards all tested insecticides varied considerably (Table 2). The maximum LD50 value of pirimicarb was 3679 times higher than the minimum LD50 of imidacloprid. The median LD50 value of all pesticides was 1.21 *μ*g a.i./bee. About 69% of substances had LD50 values below 2 *μ*g a.i./bee whereas 38% had LD50s under 0.2 *μ*g a.i./bee. Bee fresh-weight differed in all tests (range 77.7 to 112.7 mg). This led to deviations of maximum 23% (indoxacarb) and 15% (thiacloprid) from mean weight bees when calculating weight-normalised LD50 values.

**Table 2.**
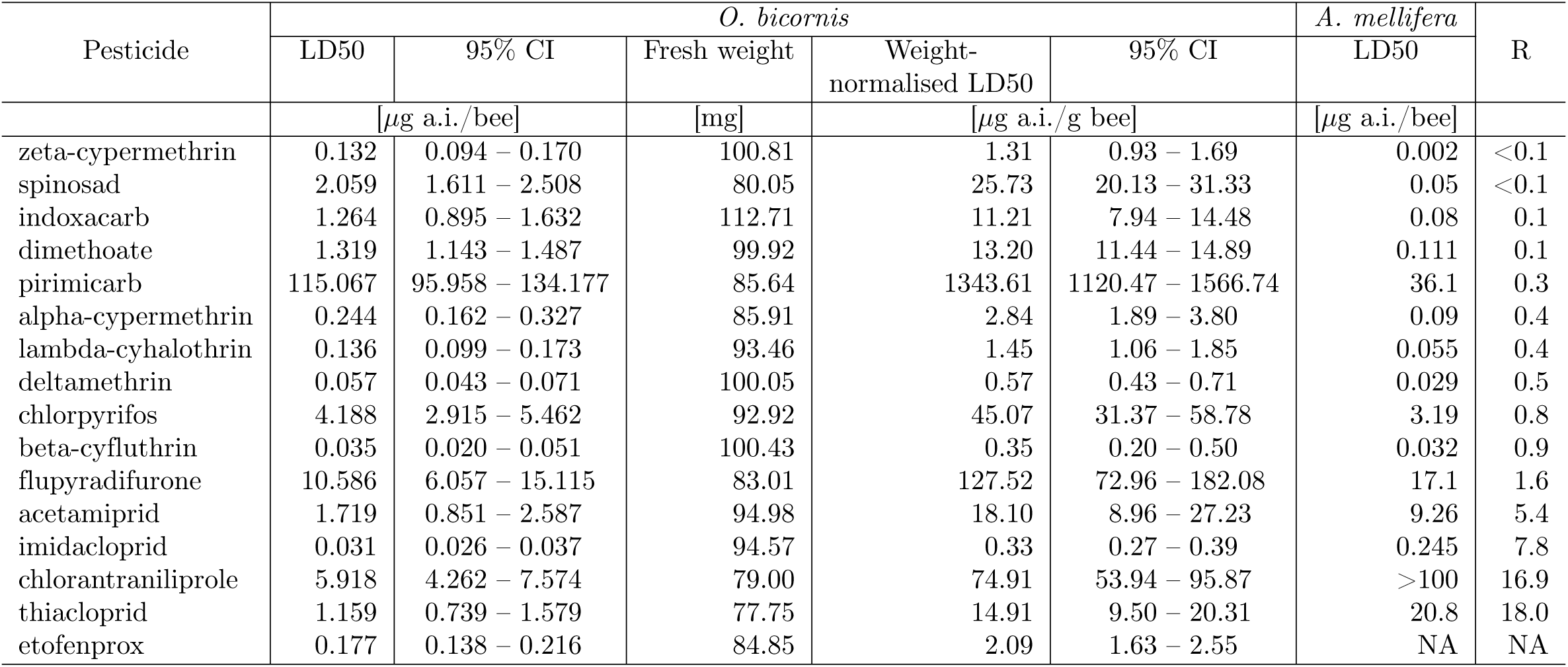
Comparison of *O. bicornis* acute toxicity endpoints from tests with honey bee regulatory endpoints. Insecticides are sorted by sensitivity ratio.

In two thirds of all cases *O. bicornis* was less sensitive than the honey bee (15 out of 16 insecticides could be evaluated). When applying an assessment factor of 10 on the respective honey bee endpoint, it was lower than the *O. bicornis* endpoint for 87% of all tested substances (Table 2). The two remaining insecticides where *O. bicornis* would still be more sensitive are formulations of chlorantraniliprole and thiacloprid. When analysing sensitivity ratios by insecticide class it was shown that for organophosphates and pyrethroids values are all below one, i.e. *O. bicornis* was more resitant than the honey bee (Fig. 1). In the case of the three tested neonicotinoids *O. bicornis* was always more sensitive.

**Fig 1.**
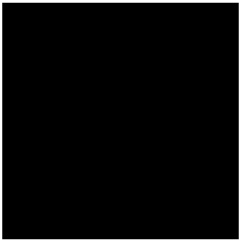
Sensitivity ratio (R) of all tested insecticides grouped by insecticide class. The dotted, black line signifies equal sensitivity of *O. bicornis* and *A. mellifera.* The dashed, red line indicates the proportion of species that would be protected when applying an assessment factor of 10 on the honey bee endpoint. The violin plot on the right shows the distribution of data points.

## Discussion

In our study we assessed the acute contact toxicity of several insecticides from several classes towards *O. bicornis.* Our goal was to compare these data to honey bee endpoints obtained from the pesticide registration process to infer on the suitability of *O. bicornis* as an additional regulatory surrogate species. Furthermore, we wanted to infer if applying an assessment factor of 10 on honey bee LD50 values would be protective for *O. bicornis*.

Acute sensitivity of *O. bicornis* varied substantially between pesticides which was expected given that the available honey bee endpoints also vary considerably (Table 2). Mean *O. bicornis* female weight also fluctuated between tests which might have slightly affected their measured sensitivity. However, this effect was not big enough to affect the toxic order of insecticides. Therefore, these LD50 values are still valid for the comparison with regulatory honey bee values. Since bee individual weight is one factor that influences sensitivity towards pesticides [16], calculating toxicity on a per weight basis leads to more precise and comparable results. Consequently, acute toxicity endpoints should generally also be reported in a weight-normalised format (see Table 2).

To create a more protective environmental risk assessment for bees, EFSA (2013) proposed the inclusion of two additional wild bee species as surrogates (*B. terrestris*, *O. bicornis*) [15]. These species should accompany the previous sole test species, the honey bee. However, in acute toxicity testing the addition of new species is only reasonable if they are generally more sensitive than the test species already in place. For two thirds of the insecticides we tested *O. bicornis* was indeed less sensitive than the honey bee (Table 2). This trend is in agreement with the findings of Uhl et al. (2016) who performed acute contact toxicity tests with five bee species and combined their dataset with LD50 values taken from literature [16]. They found that both proposed test species, *O. bicornis* and *B. terrestris*, were less sensitive towards dimethoate than several bee species, including the honey bee. Heard et al. (2017) conducted acute to chronic oral tests (up to 240h) with *B. terrestris* and *O. bicornis* and four organic pesticides, cadmium and arsenic [17]. Their results were inconclusive as to whether the wild bee species or the honey bee was acutely more sensitive. However, they could show that both newly proposed test species were less sensitive in 40% of comparisons across time. When evaluating this combined information it becomes evident that *O. bicornis* (and possibly *B. terrestris*) is seldomly an adequate supplementary surrogate species for acute testing of pesticides since its inclusion would not provide additional safety for the risk assessment process for most pesticides. As postulated by Uhl et al. (2016), test species should be chosen according to their sensitivity in acute effect studies [16]. However, both proposed test species were selected because they are bred for commercially pollination, can therefore be obtained easily in large numbers and can cope well with laboratory conditions. While those criteria are important for the conduction of laboratory experiments in general, they should not be decisive for the selection of surrogate species. The honey bee may be a better choice in acute toxicity tests since the not fully matured cuticle of young workers makes it more susceptible towards pesticides compared to solitary bees [25, 26]. Furthermore, there are differences in the immune response of young adults. In honey bees the individual detoxification capacity is relatively low and increases from thereon as they age [27, 28]. However, antioxidant enzyme levels already rise in *O. bicornis* adults before eclosion which is further evidence that they are more resistant than honey bees at least at this life stage [26].

Furthermore, we could show that for 87% of the tested insecticides an assessment factor of 10 when applied to the honey bee endpoint is sufficient to cover *O. bicornis*’ sensitivity (Fig. 1). This assessment factor was found to be protective in 95% of all cases that were analysed in the meta-analysis of Arena & Sgolastra (2014) [14]. After testing multiple wild bee species with dimethoate, Uhl et al. (2016) reaffirmed this result using a species sensitivity distribution (SSD) approach [16]. Moreover, Heard et al. (2017) also state that the honey bee is an adequate surrogate species for acute testing as long as a reasonable assessment factor is applied [17]. However, they also note that there are exceptions for some substances, e.g. neonicotinoids. Arena & Sgolastra (2014) already mentioned that for this class wild bee species showed equal to higher sensitivity than the honey bee [14]. This trend is also visible in our data: *O. bicornis* was more sensitive towards all three tested neonicotinoids (acetamiprid, imidacloprid, thiacloprid) than the honey bee (Fig. 1).

Consequently, the honey bee is a sufficient surrogate species to assess acute toxicity of most pesticides. In some cases (e.g. neonicotinoids) it might be necessary to increase the assessment factor to >10 to achieve a proper level of safety. To distinguish these substances that are relatively more harmful to wild bees than to honey bees, a comprehensive ecotoxicological database should be established that includes a representative amount of species and pesticides. Such a database would also be helpful for choosing suitable additional test species if necessary. Moreover, regulatory reporting standards should be improved. Our search for honey bee endpoints that were used in the registration process presented quite complicated. We partly received contrasting information from several sources. A solution for this problem would be the creation of a transparent and publicly available database of regulatory data. Those data could be then complemented by non-regulatory study results to further not only the open science idea but also establish a more transparent regulation process.

Despite only rarely providing additional safety for lower tier risk assessment it should be noted that both proposed test species may be more valuable surrogates in more realistic experimental setups in higher tier risk assessment. Due to their ecological differences to the honey bee, populations of *O. bicornis* and *B. terrestris* may react quite differently in (semi-)field studies. Such divergent effects have been shown in a Swedish field study where clothianidin/beta-cyfluthrin treatment of oilseed rape had no adverse effects on honey bee colonies, yet substantial impact on *O. bicornis*’ and *B. terrestris*’ population development [29]. Due to their properties they are good representatives to measure the ecological impact of pesticides on solitary bee and bumble bee species, respectively, in large field studies such as Peters et al. (2016) and Sterk et al. (2016) [30, 31].

## Conclusion

For the majority of substances we tested, the honey bee was more sensitive than *O. bicornis*. We therefore agree with Heard et al. (2017) that *A. mellifera* is a sufficient proxy for wild bee species in laboratory acute mortality testing [17]. However, it is still necessary to investigate less well-known issues such as effects of pesticides mixtures [32, 33], prolonged pesticides exposure [17] or effects of pesticide adjuvants [34]. An assessment factor of 10 proved to be protective for *O. bicornis* when applied honey bee endpoints for nearly all tested insecticides. There might be exceptions (e.g. neonicotinoids) where this assessment factor needs to be increased. Therefore, our study provides further evidence that *O. bicornis* is rarely an adequate surrogate species that will usually not improve lower tier risk assessment. Unnecessary acute studies with non-sensitive species should not be conducted. Only sensitive species should be chosen as additional surrogates to reduce overall uncertainty. However, we agree that both proposed test species can be very relevant in higher tier risk assessment. In complex field settings ecological differences between the honey bee, bumble bees and solitary bees are more relevant as shown by Rundlöf et al. (2015) [29]. Therefore, such realistic experiments are better suited to evaluate the overall impact of pesticides on wild bee species. Consequently, we believe that (semi-)field data should be relied upon to a greater extent than laboratory results in wild bee risk assessment.

## Supporting information

**S1 Appendix. Data collection of regulatory honey bee endpoints.**

**S1 Table. Overview of all tested insecticides and test dates.** For a detailed account of raw data from all tests please see Uhl et al. (2018) [22].

**S2 Table. Data sources of honey bee acute endpoints for all tested insecticides.**

**S3 Table. Different organisations that aided with data collection and contact at the respective institutions.**

## Acknowledgments

We would like to thank Therese Bürgi for her help with every laboratory-related issue. Further thanks are in order to Claudia Wollmann who performed one test with dimethoate as part of her master thesis. We are grateful to the German Environment Agency (UBA), the German Federal Office of Consumer Protection and Food Safety (BVL), EFSA, Bayer Crop Science, Dow AgroSciences and Syngenta for providing regulatory data and aiding in the data collection and validation process. Moreover, we appreciate that Syngenta, DuPont (now DowDuPont) and Belchim Crop Protection sent us samples of insecticides for testing.

## Author contributions

**Conceptualisation** PU CAB.

**Data curation** PU.

**Formal analysis** PU RSS OA.

**Funding acquisition** PU CAB.

**Investigation** RSS OA PU.

**Methodology** PU RSS OA.

**Project administration** PU CAB.

**Resources** PU RSS OA CAB.

**Software** PU RSS OA.

**Supervision** PU CAB.

**Validation** PU.

**Visualisation** PU.

**Writing - original draft** PU.

**Writing - review & editing** PU RSS OA CAB.

---

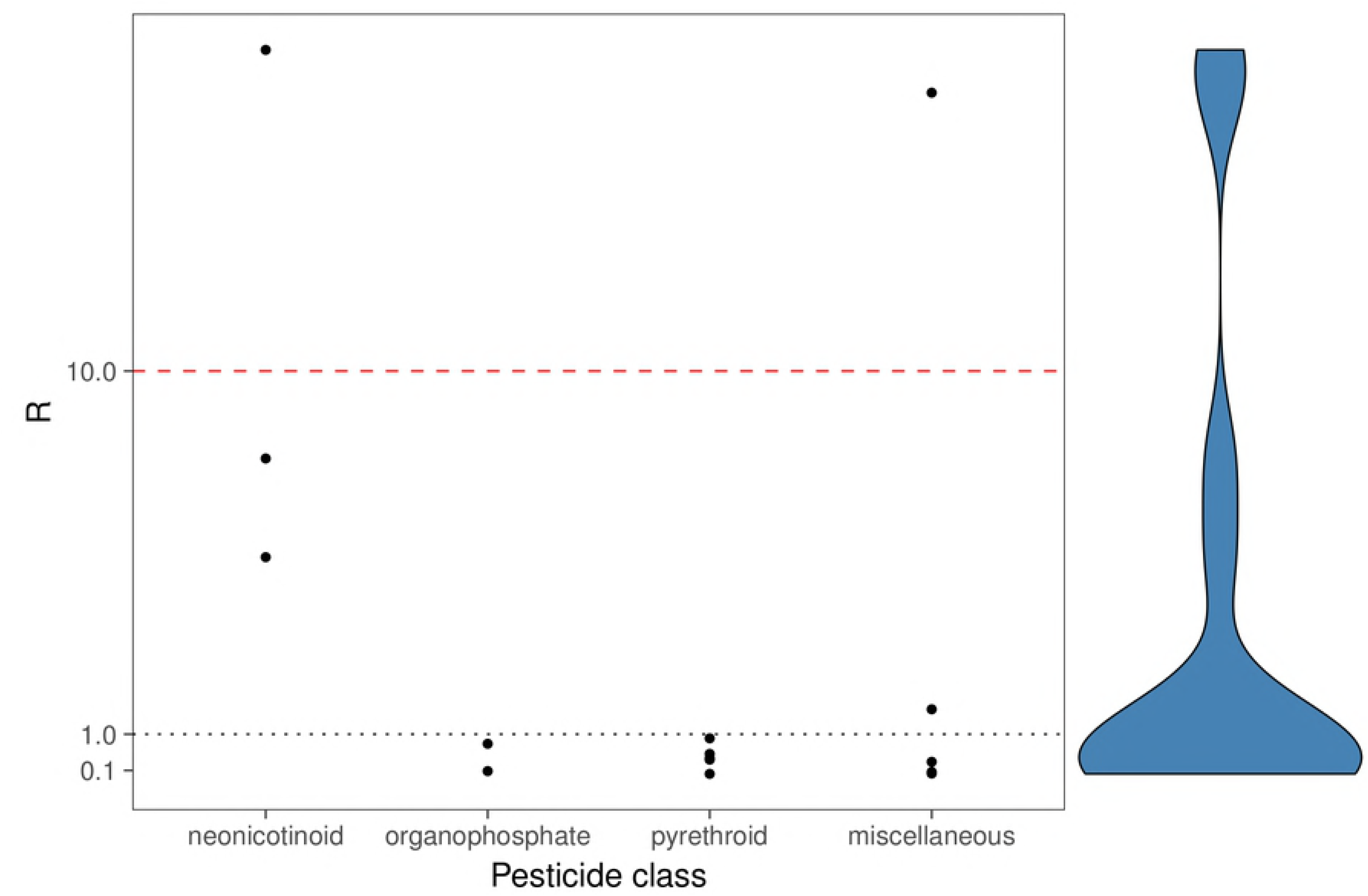

